# Novel blood pressure locus and gene discovery using GWAS and expression datasets from blood and the kidney

**DOI:** 10.1101/110833

**Authors:** Louise V. Wain, Ahmad Vaez, Rick Jansen, Roby Joehanes, Peter J. van der Most, A. Mesut Erzurumluoglu, Paul O'Reilly, Claudia P. Cabrera, Helen R. Warren, Lynda M. Rose, Germaine C. Verwoert, Jouke-Jan Hottenga, Rona J. Strawbridge, Tonu Esko, Dan E. Arking, Shih-Jen Hwang, Xiuqing Guo, Zoltan Kutalik, Stella Trompet, Nick Shrine, Alexander Teumer, Janina S. Ried, Joshua C. Bis, Albert V. Smith, Najaf Amin, Ilja M. Nolte, Leo-Pekka Lyytikäinen, Anubha Mahajan, Nicholas J. Wareham, Edith Hofer, Peter K. Joshi, Kati Kristiansson, Michela Traglia, Aki S. Havulinna, Anuj Goel, Mike A. Nalls, Siim Sõber, Dragana Vuckovic, Jian'an Luan, Fabiola Del Greco M., Kristin L. Ayers, Jaume Marrugat, Daniela Ruggiero, Lorna M. Lopez, Teemu Niiranen, Stefan Enroth, Anne U. Jackson, Christopher P. Nelson, Jennifer E. Huffman, Weihua Zhang, Jonathan Marten, Ilaria Gandin, Sarah E Harris, Tatijana Zemonik, Yingchang Lu, Evangelos Evangelou, Nabi Shah, Martin H. de Borst, Massimo Mangino, Bram P. Prins, Archie Campbell, Ruifang Li-Gao, Ganesh Chauhan, Christopher Oldmeadow, Gonçalo Abecasis, Maryam Abedi, Caterina M. Barbieri, Michael R. Barnes, Chiara Batini, John Beilby, BIOS Consortium, Tineka Blake, Michael Boehnke, Erwin P. Bottinger, Peter S. Braund, Morris Brown, Marco Brumat, Harry Campbell, John C. Chambers, Massimiliano Cocca, Francis Collins, John Connell, Heather J. Cordell, Jeffrey J. Damman, Gail Davies, Eco J. de Geus, Renée de Mutsert, Joris Deelen, Yusuf Demirkale, Alex S.F. Doney, Marcus Dörr, Martin Farrall, Teresa Ferreira, Mattias Frånberg, He Gao, Vilmantas Giedraitis, Christian Gieger, Franco Giulianini, Alan J. Gow, Anders Hamsten, Tamara B. Harris, Albert Hofman, Elizabeth G. Holliday, Jennie Hui, Marjo-Riitta Jarvelin, Åsa Johansson, Andrew D. Johnson, Pekka Jousilahti, Antti Jula, Mika Kähönen, Sekar Kathiresan, Kay-Tee Khaw, Ivana Kolcic, Seppo Koskinen, Claudia Langenberg, Marty Larson, Lenore J. Launer, Benjamin Lehne, David C.M. Liewald, Lifelines Cohort Study, Li Lin, Lars Lind, François Mach, Chrysovalanto Mamasoula, Cristina Menni, Borbala Mifsud, Yuri Milaneschi, Anna Morgan, Andrew D. Morris, Alanna C. Morrison, Peter J. Munson, Priyanka Nandakumar, Quang Tri Nguyen, Teresa Nutile, Albertine J. Oldehinkel, Ben A. Oostra, Elin Org, Sandosh Padmanabhan, Aarno Palotie, Guillaume Paré, Alison Pattie, Brenda W.J.H. Penninx, Neil Poulter, Peter P. Pramstaller, Olli T. Raitakari, Meixia Ren, Kenneth Rice, Paul M. Ridker, Harriëtte Riese, Samuli Ripatti, Antonietta Robino, Jerome I. Rotter, Igor Rudan, Yasaman Saba, Aude Saint Pierre, Cinzia F. Sala, Antti-Pekka Sarin, Reinhold Schmidt, Rodney Scott, Marc A. Seelen, Denis C. Shields, David Siscovick, Rossella Sorice, Alice Stanton, David J. Stott, Johan Sundström, Morris Swertz, Kent D. Taylor, Simon Thom, Ioanna Tzoulaki, Christophe Tzourio, André G. Uitterlinden, Understanding Society Scientific group, Uwe Vöker, Peter Vollenweider, Sarah Wild, Gonneke Willemsen, Alan F. Wright, Jie Yao, Sébastien Thériault, David Conen, Attia John, Peter Sever, Stéphanie Debette, Dennis O. Mook-Kanamori, Eleftheria Zeggini, Tim D. Spector, Pim van der Harst, Colin N.A. Palmer, Anne-Claire Vergnaud, Ruth J.F. Loos, Ozren Polasek, John M. Starr, Giorgia Girotto, Caroline Hayward, Jaspal S. Kooner, Cecila M. Lindgren, Veronique Vitart, Nilesh J. Samani, Jaakko Tuomilehto, Ulf Gyllensten, Paul Knekt, Ian J. Deary, Marina Ciullo, Roberto Elosua, Bernard D. Keavney, Andrew A. Hicks, Robert A. Scott, Paolo Gasparini, Maris Laan, YongMei Liu, Hugh Watkins, Catharina A. Hartman, Veikko Salomaa, Daniela Toniolo, Markus Perola, James F. Wilson, Helena Schmidt, Jing Hua Zhao, Terho Lehtimäki, Cornelia M. van Duijn, Vilmundur Gudnason, Bruce M. Psaty, Annette Peters, Rainer Rettig, Alan James, J Wouter Jukema, David P. Strachan, Walter Palmas, Andres Metspalu, Erik Ingelsson, Dorret I. Boomsma, Oscar H. Franco, Murielle Bochud, Christopher Newton-Cheh, Patricia B. Munroe, Paul Elliott, Daniel I. Chasman, Aravinda Chakravarti, Joanne Knight, Andrew P. Morris, Daniel Levy, Martin D. Tobin, Harold Snieder, Mark J. Caulfield, Georg B. Ehret

## Abstract

Elevated blood pressure is a major risk factor for cardiovascular disease and has a substantial genetic contribution. Genetic variation influencing blood pressure has the potential to identify new pharmacological targets for the treatment of hypertension. To discover additional novel blood pressure loci, we used 1000 Genomes Project-based imputation in 150,134 European ancestry individuals and sought significant evidence for independent replication in a further 228,245 individuals. We report 6 new signals of association in or near *HSPB7, TNXB, LRP12, LOC283335, SEPT9* and *AKT2*, and provide new replication evidence for a further 2 signals in *EBF2* and *NFKBIA*. Combining large whole-blood gene expression resources totaling 12,607 individuals, we investigated all novel and previously reported signals and identified 48 genes with evidence for involvement in BP regulation that are significant in multiple resources. Three novel kidney-specific signals were also detected. These robustly implicated genes may provide new leads for therapeutic innovation.

## INTRODUCTION

Genetic support for a drug target increases the likelihood of success in drug development (1) and there is clear unmet need for novel therapeutic strategies to treat individuals with hypertension (2). A number of large studies have described blood pressure (BP) variant identification by genome-wide and targeted association approaches (3-19). Clinically the most predictive BP traits for cardiovascular risk are systolic blood pressure (SBP) and diastolic blood pressure (DBP), reflecting roughly the peak and trough of the BP curve, and pulse pressure (PP), the difference between SBP and DBP (20) reflecting arterial stiffness. Using these three traits, we undertook a meta-analysis of 150,134 individuals from 54 genome-wide association studies of European ancestry with imputation based on the 1000 Genomes Project Phase 1. To minimize reporting of false positive associations, we sought stringent evidence for significant independent replication in a further 228,245 individuals. We further followed up novel and previously reported association signals in multiple large gene expression databases and the largest kidney tissue gene expression resource currently available. Finally, we searched for enrichment of associated genes in biological pathways and gene sets and identified whether any of the genes were known drug targets.

## RESULTS

The stage 1 discovery meta-analysis included 150,134 individuals (**Online Methods; Supplementary Tables 1-4, Supplementary Figures 1** and **2**) and 7,994,604 variants with minor allele frequency (MAF) >1% and an effective sample size of at least 60% of the total (**Online Methods**). We identified 61 signals in the discovery analysis that were candidates for novel BP signals (*P* < 10^−6^ for any trait; **Supplementary Table 5**). To ensure robustness of signals, we examined BP associations in an additional 228,245 individuals from 15 independent studies for replication, including 140,886 individuals from UK Biobank (19) (**Supplementary Table 6** and **Online Methods**). We used the most significant (“sentinel”) SNP and trait for each locus in replication (61 tests). Twenty-two putatively novel association signals were initially confirmed showing significant evidence of replication in the independent stage-2 studies (P < 8.2x10^−4^, Bonferroni correction for 61 tests) and genome-wide significance (P < 5x10^−8^) in a meta-analysis across all 378,376 individuals (**Online methods, Table 1, Supplementary Table 7**). Of these, 14 were subsequently published in two other studies (18,19) which presented genome-wide significant associations with evidence of replication. A further two were highlighted as putative novel signals in one of those studies (18) but had not been confirmed by replication. In our study, we report the 6 remaining novel signals, and the 2 previously unconfirmed signals (in *EBF2* and in *NFKBIA*), as novel signals. The 8 novel signals included 7 signals at 7 independent loci (**Supplementary Figure 3**) and one novel independent signal near a previously reported hit near *TNXB* (**Online Methods, Supplementary Table 8, Supplementary Figure 4**). The novel signals show both significant evidence of replication in the independent stage-2 studies (*P* < 8.2x10^−4^, Bonferroni correction for 61 tests) and genome-wide significance (*P<* 5x10^−8^) in a metaanalysis across all 378,376 individuals. The sentinel variants at all 8 signals were common (MAF>5%) and the novel secondary signal at *TNXB* was in high linkage-disequilibrium (*r^2^* > 0.8) with a non-synonymous SNP. With the exception of rs9710247, which was only significant for association with DBP, all signals were significantly associated (P<0.006, Bonferroni corrected for 8 tests) with all 3 traits (**Table 1** and **Supplementary Table 9**).

**Table 1:**
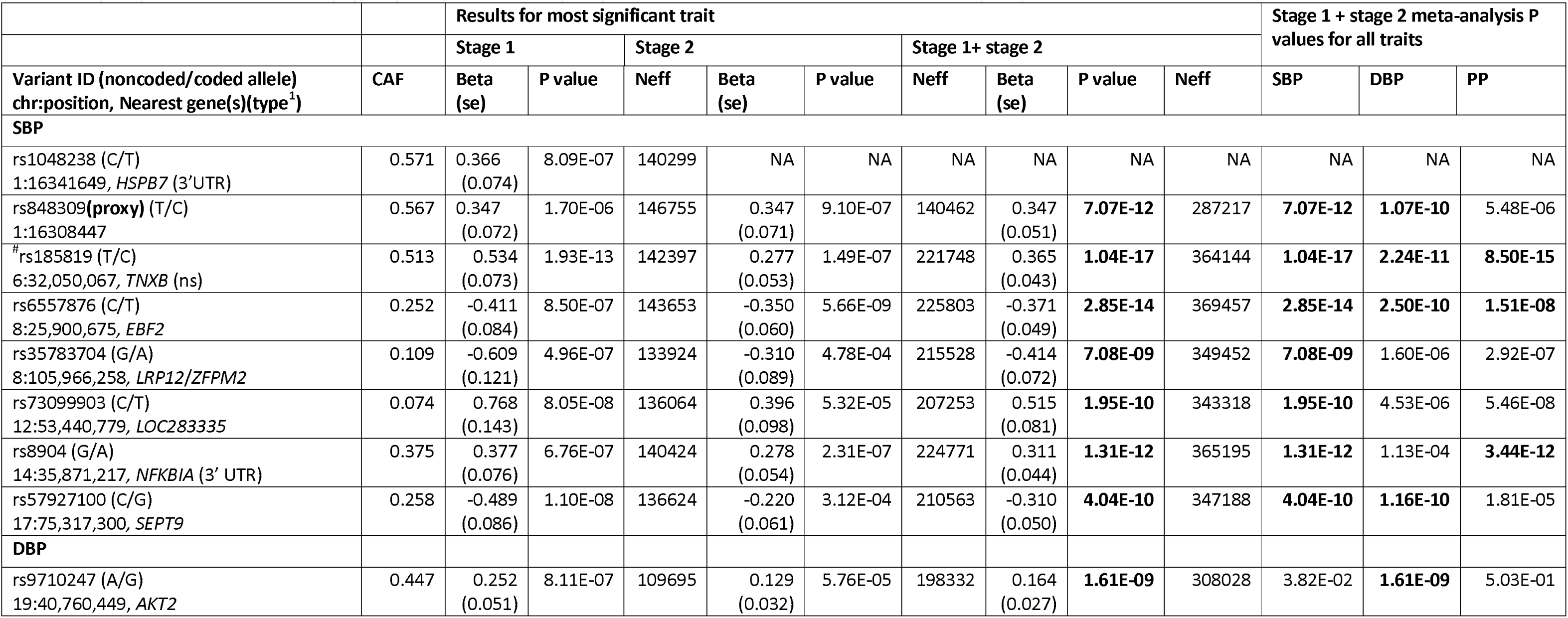
Novel genome-wide significant signals of association. Results from stage 1 and stage 2, and the meta-analysis of stage 1 and stage 2, for all novel genome-wide significant signals of association. *P* values of association for all 3 traits from a meta-analysis of stages 1 and 2 are also presented. Genome-wide significant *P* values (P<5x10^−8^) are in bold. Abbreviations: CAF: coded allele frequency se: standard error, Neff: effective sample size. ^#^Novel signal at previously reported locus. ^1^For intragenic variants the nearest genes are listed, all other variants are intronic unless indicated otherwise; ns= non-synonymous, s=synonymous, UTR= Untranslated Region. Results from proxy SNPs are indicated by (**proxy**); rs848309 was a proxy SNP for rs1048238 and rs10926988 was a proxy SNP for chr1:243458005:I.

We next sought to identify which genes might have expression levels that were associated with genotypes of the BP-associated variants reported in this study and others. Strong evidence of an association with expression of a specific gene may provide clues as to which gene(s) might be functionally relevant to that signal. We took the 139 BP association signals reported prior to these studies (18,19), and 22 novel signals of association identified and confirmed in this study and two contemporaneous studies (3-19, 21) (**Supplementary Table 10**), and searched for evidence of association with gene expression in whole-blood (four studies, total n=12,607; **Online Methods**) and in kidney tissue (n=134, the largest kidney eQTL resource currently available). Although of unclear direct relevance to BP, whole-blood was studied due to the availability of large data sets enabling a powerful assessment of expression patterns that are likely present across multiple cell and tissue types. Kidney was chosen because of the many renal pathways that regulate BP and outstanding questions about the relevance of kidney pathways to the genetic component of BP regulation in the general population (3,15). Expression quantitative trait loci (eQTL) signals were filtered by false discovery rate (FDR<5%) and we examined *cis* (within 1Mb) associations only (**Online methods** and **Supplementary Material**).

The four blood eQTL data sets were NESDA-NTR (22, 23), SABRe (15), the BIOS resource (24) and GTEx(25) (**Online Methods and Supplementary Material**). The BIOS resource (n=2,116) has not previously been utilized in the analysis of BP associations, findings from NESDA-NTR and SABRe have been reported for a subset of the previously published signals (16,17). For a total of 369 genes, gene-expression was associated with the BP SNP in one or more of the 4 blood datasets at experiment-wide significance (**Supplementary Table 11**). This included 14 genes for 6 of the 8 novel signals. For 110 genes, we found eQTL evidence in 2 out of 4 datasets (**Figure 1**), including 4 genes for 2 of the novel signals; *EIF4B* and *TNS2* for rs73099903 and *MAP3K10* and *PLD3* for rs9710247. SNP rs73099903 was in strong linkage disequilibrium (LD r^2^>0.9) with the SNP most strongly associated with *TNS2* expression in the BIOS resource. *TNS2* encodes a tensin focal adhesion molecule and may have a role in renal function (26).

**Figure 1:**
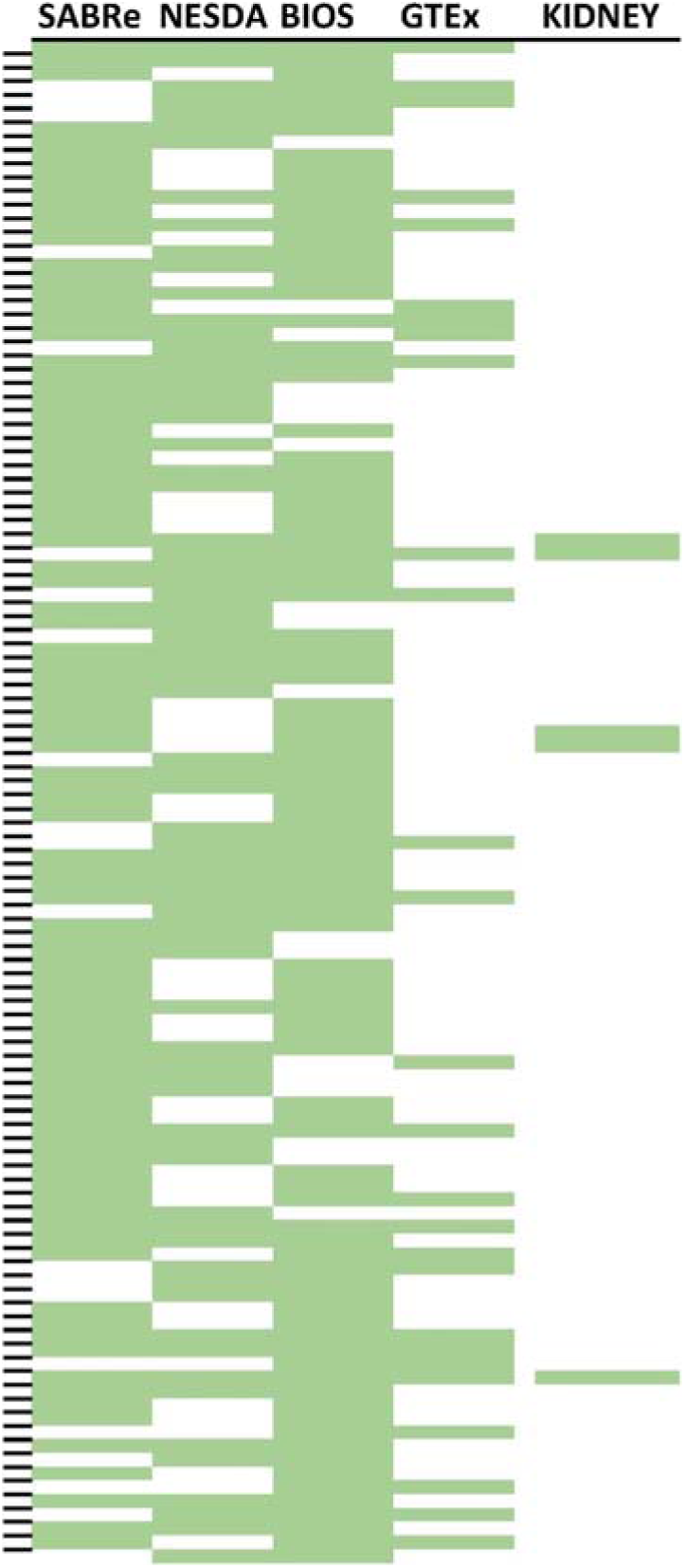
Overlap of eQTL evidence from four whole-blood and one kidney resource. The figure indicates overlap of evidence for eQTLs from four whole-blood studies (SABRe, NESDA-NTR, BIOS, and GTEx) and from one kidney resource (TransplantLines). Every colored line indicates that this gene was analysis-wide significant in a given resource (see **Online Methods**). Only genes identified by at least two resources are shown. The genes are sorted by genomic position on the y-axis.

For 48 genes, we found evidence in 3 out of the 4 resources (**Table 2**), suggesting robustness of the SNP-gene expression correlation signal and highlighting those genes as potential candidates in genetic BP regulation. Of the 48 genes, 28 have not previously been described in eQTL analyses using BP associated SNPs and all were correlated with previously reported BP association signals.

**Table 2:** BP associated SNPs associated with expression of the same gene across 4 or 3 independent whole-blood eQTL resources and the kidney resource. Signals of association of SNP genotype and gene expression in other non-blood tissues in GTEx and in kidney are also indicated. Blood dataset order: (i) SABRe, (ii) NESDA-NTR, (iii) BIOS, (iv) GTEx (whole-blood). Top eQTL: Top GWAS SNP is top eQTL SNP (or in high LD, *r^2^*>0.9, with top eQTL SNP) in at least one data set. eQTL signal previously reported: Genes for which eQTL signals have been previously reported for that sentinel SNP(15-17). For full list, see **Supplementary Table 12.**

| Sentinel SNP | Chr | Position | Gene | Blood data sets | Top eQTL | Signal in other tissue(s) in GTEx | Signal in kidney | eQTL signal previously reported |
| --- | --- | --- | --- | --- | --- | --- | --- | --- |
| **Signal in 4 whole-blood eQTL resources** |  |  |  |  |  |  |  |  |
| rs17367504 | 1 | 11862778 | *CLCN6* | YYYY |  | Y |  | Y |
| rs2169137 | 1 | 204497913 | *MDM4* | YYYY | Y | Y |  | Y |
| rs10926988 | 1 | 243483279 | *SDCCAG8* | YYYY |  | Y |  |  |
| rs319690 | 3 | 47927484 | *MAP4* | YYYY | Y | Y |  | Y |
| rs12521868 | 5 | 131784393 | *SLC22A5* | YYYY |  | Y |  |  |
| rs900145 | 11 | 13293905 | *ARNTL* | YYYY |  | Y |  | Y |
| rs1060105 | 12 | 123806219 | *CDK2AP1* | YYYY | Y | Y | Y |  |
| rs1378942 | 15 | 75077367 | *SCAMP2* | YYYY |  |  |  |  |
| rs1126464 | 16 | 89704365 | *CHMP1A* | YYYY |  | Y |  | Y |
| rs1126464 | 16 | 89704365 | *FANCA* | YYYY |  |  |  | Y |
| rs12946454 | 17 | 43208121 | *DCAKD* | YYYY |  | Y | Y | Y |
| **Signal in 3 (out of 4) whole-blood eQTL resources** |  |  |  |  |  |  |  |  |
| rs17367504 | 1 | 11862778 | *MTHFR* | YYYN |  | Y |  | Y |
| rs871524 | 1 | 38411445 | *FHL3* | NYYY |  | Y |  |  |
| rs871524 | 1 | 38411445 | *SF3A3* | NYYY |  | Y |  |  |
| rs4660293 | 1 | 40028180 | *PABPC4* | YYYN | Y | Y |  | Y |
| rs6749447 | 2 | 169041386 | *STK39* | YYYN | Y |  |  |  |
| rs347591 | 3 | 11290122 | *ATG7* | YYYN |  | Y |  |  |
| rs319690 | 3 | 47927484 | *ZNF589* | YYNY |  | Y |  |  |
| rs12521868 | 5 | 131784393 | *SLC22A4* | YYYN |  | Y |  |  |
| rs1563788 | 6 | 43308363 | *CRIP3* | YYYN | Y |  |  | Y |
| rs10943605 | 6 | 79655477 | *PHIP* | YYYN | Y | Y |  | Y |
| rs4728142 | 7 | 128573967 | *IRF5* | NYYY |  | Y | Y | Y |
| rs4728142 | 7 | 128573967 | *TNPO3* | YYYN |  |  | Y |  |
| rs2898290 | 8 | 11433909 | *BLK* | YYYN |  | Y |  |  |
| rs2898290 | 8 | 11433909 | *FAM167A* | NYYY |  | Y |  |  |
| rs2898290 | 8 | 11433909 | *FDFT1* | YYYN |  | Y |  |  |
| rs2071518 | 8 | 120435812 | *NOV* | YYYN |  | Y |  |  |
| rs76452347 | 9 | 35906471 | *TPM2* | YYYN |  |  |  |  |
| rs10760117 | 9 | 123586737 | MEGF9 | YYYN |  | Y |  | Y |
| rs4494250 | 10 | 96563757 | HELLS | YYYN |  |  |  | Y |
| rs11191548 | 10 | 104846178 | NT5C2 | YYYN | Y |  |  |  |
| rs661348 | 11 | 1905292 | TNNT3 | NYYY |  | Y |  |  |
| rs2649044 | 11 | 9763969 | SBF2 | YYYN |  |  |  |  |
| rs2649044 | 11 | 9763969 | SWAP70 | YYYN | Y | Y |  | ? |
| rs7129220 | 11 | 10350538 | ADM | YYYN |  |  |  | Y |
| rs7103648 | 11 | 47461783 | MYBPC3 | YYYN |  |  |  |  |
| rs3741378 | 11 | 65408937 | CTSW | YYYN |  |  |  |  |
| rs7302981 | 12 | 50537815 | LIMA1 | YYYN |  |  |  | Y |
| rs7302981 | 12 | 50537815 | ATF1 | YYNY |  | Y |  |  |
| rs1036477 | 15 | 48914926 | FBN1 | YNYY |  |  |  |  |
| rs1378942 | 15 | 75077367 | CSK | YYYN | Y | Y |  | Y |
| rs1378942 | 15 | 75077367 | MPI | NYYY |  | Y |  |  |
| rs1378942 | 15 | 75077367 | ULK3 | YNYY |  | Y |  | Y |
| rs12946454 | 17 | 43208121 | NMT1 | YYYN |  |  |  | Y |
| rs2304130 | 19 | 19789528 | GATAD2A | YYYN |  |  |  |  |
| rs867186 | 20 | 33764554 | EIF6 | NYYY |  | Y |  |  |
| rs6095241 | 20 | 47308798 | PREX1 | YYYN |  |  |  |  |
| rs9306160 | 21 | 45107562 | RRP1B | YNYY | Y | Y |  |  |

In the kidney dataset (TransplantLines) (27), there was association of gene expression and genotype for nine SNPs and 13 genes **(Table 2, Figure 1** and **Supplementary Table 12**). Nine of the SNP-gene expression associations were also observed in the whole-blood eQTL datasets, suggesting that those signals may not be unique to the kidney. We report three signals that were unique to the kidney and not previously reported (*C4orf34, HIP2* and *ASIC1*) and confirm a previously reported kidney eQTL signal for an anti-sense RNA for PSMD5 (15). The same SNP was also an eQTL for *PSMD5* itself in both blood and kidney. *ASIC1* encodes the Acid Sensing Ion Channel Subunit 1 which may interact (and be co-expressed) with ENaC subunits which mediate trans-epithelial Na transport in the kidney (28). The comparatively small number of signals using kidney tissue (**Table 2** and **Figure 1**) compared to whole-blood could be due to the small sample size.

For genes implicated by eQTL information from whole-blood, we tested for enrichment of biological pathways and gene ontologies (**Online Methods**). We noted enrichment of the 48 genes implicated by 3 or 4 blood eQTL resources, **Table 2**, and a further 53 genes containing a non-synonymous variant with *r*^2^ > 0.5 with the top SNP (**Supplementary Table 13**), in pathways and ontology terms related to actin and striated muscle (**Supplementary Tables 14 and 15, Online Methods**). Network analysis using the same genes highlighted further GO terms relating to muscle function, particularly cardiac muscle (**Online Methods, Supplementary Table 16**). We tested the overlap of 161 non-HLA BP associated variants with DNase Hypersensitivity sites identified in the Roadmap and ENCODE cell lines (**Online Methods**) and identified an overall enrichment in multiple cell and tissue types including heart, kidney and smooth muscle (**Supplementary Figure 5**).

We next investigated these genes for potential suitable drug targets using the drug gene interaction database (DGIdb) (29) and found 19 genes with known drug-gene interactions and 17 additional genes with predicted druggability (**Supplementary Table 17**). These findings highlight potential opportunities for novel therapeutic development and possible drug re-purposing, given that a large number of the genes is already now targetable.

## DISCUSSION

Enhanced discovery of BP loci increases the potential targets for therapeutic advances. After major advances in the number of BP loci known over the last years and months, we report 8 novel signals that implicate 5 regions of the genome not previously connected to blood pressure regulation.

Six of the 8 novel signals we report had not previously been reported. Two signals (in *EBF2* and *NFKBIA*) have been suggested previously but without evidence for replication (18). For these two signals we present, for the first time, stringent evidence of replication, confirming their relevance to blood pressure genetics.

The path from signal to genes is the essential next step towards realizing the therapeutic potential of a genetic locus and understanding the mechanisms of BP regulation. We have used several large eQTL resources as a first step to realize this objective. As expected, we observed that even across eQTL studies of the same tissue, there is limited overlap in experiment-wide significant signals suggesting either biologic variability, technology-specific differences in coverage of genes, or the possibility of false positive results despite stringent within-experiment significance thresholds. By selecting genes only significant in at least three resources, we identified 48 genes as candidates for further study. These results are limited by the availability of large eQTL resources for whole-blood only, which precludes well-powered comparisons across tissue types, particularly as the origin of blood pressure control is unlikely to be located in the blood. Enrichment and pathway analyses using these genes, and genes containing a correlated functional variant, highlight the potential relevance of muscular tissue and pathways, compatible with a vascular and cardiac origin of BP genetics, extending previous evidence (15). We identify a number of potential drug targets in the pathways identified, providing, together with previous results, a possible avenue for development of pharmacological interventions modulating blood pressure.

In summary, our study reports novel BP association signals and reports new candidate BP genes, contributing to the transition from variants to genes to explain BP variation.

## MATERIALS AND METHODS

### Studies Stage 1

Results from 54 independent European-ancestry studies, totaling 150,134 individuals, were included in the Stage 1 meta-analysis: AGES (n=3215), ARIC (n=9402), ASPS (n=828), B58C (n=6458), BHS (n=4492), CHS (n=3254), Cilento study (n=999), COLAUS (n=5404), COROGENE-CTRL (n=1878), CROATIA-Vis (n=945), CROATIA-Split (n=494), CROATIA-Korcula (n=867), EGCUT (n=6395), EGCUT2 (n=1844), EPIC (n=2100), ERF (n=2617), Fenland (n=1357), FHS (n=8096), FINRISK-ctrl (n=861), FINRISK CASE (n=839), FUSION (n=1045), GRAPHIC (n=1010), H2000-CTRL (n=1078), HealthABC (n=1661), HTO (n=1000), INGI-CARL (n=456), INGI-FVG (n=746), INGI-VB (n=1775), IPM (n=300), KORAS3 (n=1590), KORAS4 (n=3748), LBC1921 (n=376), LBC1936 (n=800), LOLIPOP-EW610 (n=927), MESA (n=2678), MICROS (n=1148), MIGEN (n=1214), NESDA (n=2336), NSPHS (n=1005), NTR (n=1490), PHASE (n=4535), PIVUS (n=945), PROCARDIS (n=1652), SHIP (n=4068), ULSAM (n=1114), WGHS (n=23049), YFS (n=1987), ORCADES (n=1908), RS1 (n=5645), RS2 (n=2152), RS3 (n=3018), TRAILS (n=1262), TRAILS-CC (n=282) and TWINGENE (n=9789). Full study names and general study information is given in **Supplementary Table 1.**

### Study-level genotyping and association testing

Three quantitative BP traits were analyzed: SBP, DBP, and PP (difference between SBP and DBP). Within each study, individuals known to be taking anti-hypertensive medication had 15 mmHg added to their raw SBP value and 10 mmHg added to their raw DBP values (30). A summary of BP phenotypes in each study is given in **Supplementary Table 2.** Association testing was undertaken according to a central analysis plan that specified the use of sex, age, age^2^, and body mass index (BMI) as covariates and optional inclusion of additional covariates to account for population stratification (**Supplementary Table 3**). Trait residuals were calculated for each trait using a normal linear regression of the medication-adjusted trait values (mmHg) onto all covariates. The genotyping array, pre-imputation quality control filters, imputation software and association testing software used by each study are listed in **Supplementary Table 4.** All studies imputed to the 1000 Genomes Project Phase 1 integrated release version 3 [March 2012] all ancestry reference panel (31). Imputed genotype dosages were used to take into account uncertainty in the imputation. Association testing was carried out using linear regression of the trait residuals onto genotype dosages under an additive genetic model. Methods to account for relatedness within a study were used where appropriate (**Supplementary Table 3**). Results for all variants (SNPs and INDELs) were then returned to the central analysis group for further quality control checks and meta-analysis.

### Stage 1 meta-analysis

Central quality control checks were undertaken across all results sets. This included checks to ensure allele frequency consistency (across studies and with reference populations), checks of effect size and standard error distributions (i.e. to highlight phenotype issues) and generation of quantile-quantile (QQ) plots and genomic inflation factor lambdas to check for over- or under-inflation of test statistics. Genomic control was applied (if lambda>1) at study-level. Variants with imputation quality <0.3 were excluded prior to meta-analysis. Inverse variance weighted meta-analysis was undertaken. After meta-analysis, variants with a weighted minor allele frequency of less than 1 % or N effective (product of study sample size and imputation quality summed across contributing studies) <60% were then excluded and meta-analysis genomic control lambda calculated and used to adjust the meta-analysis results.

### Selection of regions for follow-up

For each trait, regions of association were selected by ranking variants by *P* value, recording the variant with the lowest *P* value as a sentinel variant and then excluding all variants +/-500kb from the sentinel and re-ranking the remaining variants. This was undertaken iteratively until all sentinel variants representing 1Mb regions containing associations with *P* <10^−6^ had been identified. To identify additional signals represented by secondary sentinel variants within 500kb of each of the sentinel variants, GCTA (32) was used to run conditional analyses (conditioned on the first sentinel variant) on each of the 1Mb regions using GWAS summary statistics and LD information from ARIC. This was done both for putatively novel regions and for regions that had previously been reported. A chi-squared test of heterogeneity of effect sizes across the 54 studies was run for each sentinel variant and those with *P* <0.05 for heterogeneity were excluded from further follow-up. Variants with *P*<10^−6^ after conditioning on the sentinel SNP (novel or known) in the region and for which any attenuation of the –log10 *P* value was less than 1.5 fold, were also taken forward for replication.

### Studies stage 2

Data from 14 independent studies, totaling 87,360 individuals, and the first release of UK Biobank, totaling 140,886 individuals, were combined to replicate the findings from stage 1 (i.e. totaling 228,245 individuals). Stage 2 study details, including full study names, are given in Supplementary **Table 6** and included 3C-Dijon (n=4061), Airwave (n=14023), ASCOT-SC (n=2462), ASCOT-UK (n=3803), BRIGHT (n=1791), GAPP (n=1685), GoDARTs (n=7413), GS:SFHS (n=9749), HCS (n=2112), JUPITER (n=8718), LifeLines (n=13376), NEO (n=5731), TwinsUK (n=4973), UK Biobank-CMC (n=140,886) and UKHLS (n=7462). Analysis was undertaken using the same methods as described for Stage 1 studies. UK Biobank-CMC utilized a newer imputation reference panel than the other studies and where a requested variant was not available, a proxy was used (next most significant *P* value with linkage disequilibrium *r*^2^> 0.6 with original top variant). Results from all stage 2 studies were meta-analyzed using inverse-variance weighted meta-analysis. Two of the variants, rs1048238 and chr1:243458005:I, were not available in the largest study in Stage 2 (UK Biobank-CMC) and so proxy variants were selected (based on *P* value and LD).

### Stage 1 + Stage 2 meta-analysis

Following meta-analysis of stage 1 and stage 2 results, signals with a *P* > 5x10^−8^ were excluded. Of the signals with a final *P* <5x10^−8^, support for independent replication within the stage 2 studies only was sought. Any signals which had *P* < 5x10^−8^ and evidence for independent replication in stage 2 alone, indicated by *P* < 8.2x10^−4^ (Bonferroni correction for 61 tests) were reported as novel signals of association with BP. Any signals which were subsequently reported by other BP GWAS that were accepted for publication during the time this analysis was ongoing, or signals for which independence from another known signal could not be established, were removed from our list of novel signals at this stage (**Supplementary Table 5**).

### Genotype and gene expression

We searched for signals of association of genotype with gene expression for the 22 signals (including 8 novel) signals described in this study (**Supplementary Table 7)** and all signals reported prior to our study (**Supplementary Table 10**) (3-17, 21) in 3 whole-blood data sets, 1 kidney data set and the GTEx multiple tissue data resource, which included whole-blood (25). We selected cis signals of association which were significant after controlling for 5% False Discovery Rate (FDR). The 3 whole-blood eQTL data sets were the NHLBI Systems Approach to Biomarker Research in Cardiovascular Disease initiative whole-blood eQTL resource (SABRe) (microarray, n=5257), NESDA-NTR (microarray, n=4896), BIOS (RNAseq, n=2116). The whole-blood data from GTEx was based on data from 338 samples. The kidney data set comprised 236 donor-kidney samples from 134 donors (27). Full details of each data set can be found in the **Supplementary Material.**

### LD lookup

The 1000 Genomes Project phase 3 release of variant calls was used (Feb. 20th, 2015), using 503 subjects of European ancestry(31). *r^2^* between the sentinel SNPs and all other bi-allelic SNPs within the corresponding 2 Mb area was calculated using the Tabix and PLINK software package (v1.07) (33, 34). Annotation was performed using the ANNOVAR software package(35).

### Gene-based pathway analysis

All genes identified in 3 or 4 of the whole-blood eQTL resources above (**Table 2**), and genes containing a non-synonymous variant with *r^2^*>0.5 with the sentinel variant (Supplementary Table 13), were tested for enrichment of biological pathways and gene ontology terms using ConsensusPathDB (36) using a FDR<5% cut-off. Enriched pathways and GO terms containing genes only implicated by a single BP-associated variant were not reported.

### Network analysis

To construct a functional association network, we combined two prioritized candidate gene sets into a single query gene set as (i) genes mapping to the non-synonymous SNPs (nsSNPs) in high LD (*r^2^*>0.5) with the corresponding sentinel BP associated SNP, and (ii) genes with eQTL evidence from 3 or 4 of the blood eQTL resources. Three sentinel SNPs (rs185819, rs926552 and rs805303) mapping to the HLA region on chromosome 6 were excluded from downstream analyses. The single query gene set was then used as input for the functional network analysis(37). We used the Cytoscape (38) software platform extended by the GeneMANIA(39) plugin (Data Version: 8/12/2014)(40). All the genes in the composite network, either from the query or the resulting gene sets, were then used for functional enrichment analysis against Gene Ontology terms (GO terms) (41) to identify the most relevant GO terms using the same plugin (40).

### DNase1 Hypersensitivity overlap enrichment across tissue and cell-types

The Functional element Overlap analysis of the Results of Genome Wide Association Study (GWAS) Experiments (Forge tool v1.1)(42) was used to test for enrichment of overlap of BP SNPs in tissues and cell lines from the Roadmap and ENCODE projects. All 164 SNPs were entered and 143 were included in the analysis. SNPs from 9 commonly used GWAS arrays were used to select background sets of SNPs for comparison and 10,000 background repetitions were run. A Z-score threshold of >=3.39 (estimated false positive rate of 0.5%) was used to declare significance.

### Drug-gene interactions

Genes used for pathway and gene ontology enrichment analyses were further investigated for potential druggable targets using the drug gene interaction database (DGIdb). The known drug-gene interactions search parameters were set investigate all 15 databases in DGIdb and include all types of interactions. The analysis performed for druggability prediction included all 9 databases exclusively inspecting expert curated data only.

## NOTE

Supplementary Information and Source Data files are available in the online version of the paper.

## ACKNOWLEDGEMENTS

We thank all the study participants of this study for their contributions. Detailed acknowledgment of funding sources is provided in the **Supplementary Material.**

## CONFLICTS OF INTERESTS

The authors declare competing financial interests (see corresponding section in the Supplementary Material).

## AUTHOR CONTRIBUTIONS

### Secondary analyses

Design of secondary analyses: L.V.W., G.B.E, M.J.C., H.Snieder, M.D.T, R.Joehanes, A.V., R.Jansen, A. V., J.K., P.O.R., A.P.M., C.P.C. Computation of secondary analysis: L.V.W., G.B.E., A.P.M., M.E., T.B., L.Lin, R.Joehanes, A.V., P.J.v.d.M., R.Jansen, C.P.C.

### Discovery

WGHS: Study phenotyping: P.M.R. Genotyping or analysis: D.I.C., L.M.R. Study PI: D.I.C., P.M.R. RS: Study phenotyping: G.C.V. Genotyping or analysis: G.C.V., A.G.U. Study PI: O.H.F., A.Hofman, A.G.U.

NTR: Study phenotyping: E.J.d.G., G.W. Genotyping or analysis: J.J.H., E.J.d.G., G.W. Study PI: D.I.B., E.J.d.G.

STR: Study phenotyping: E.I. Genotyping or analysis: R.J.S., M.Frånberg Study PI: E.I., A.Hamsten

EGCUT: Genotyping or analysis: T.E. Study PI: A.Metspalu

ARIC: Genotyping or analysis: D.E.A., A.C.M., P.N. Study PI: A.Chakravarti

FHS: Study phenotyping: D.L. Genotyping or analysis: S.J.H. Study PI: D.L.

MESA: Study phenotyping: J.I.R. Genotyping or analysis: W.P., X.G., J.I.R., J.Y. Study PI: W.P.

B58C: Study phenotyping: D.P.S. Genotyping or analysis: D.P.S. Study PI: D.P.S.

COLAUS: Study phenotyping: P.V. Genotyping or analysis: M.Bochud, Z.K. Study PI: P.V.

PROSPER: Study phenotyping: J.W.J., D.J.S. Genotyping or analysis: S.Trompet, J.D. Study PI: J.W.J.

BHS: Study phenotyping: A.James Genotyping or analysis: N.Shrine, J.H., J.B.

SHIP: Study phenotyping: M.D. Genotyping or analysis: A.T., M.D., U.V. Study PI: R.R.

KORA S4: Genotyping or analysis: J.S.R. Study PI: A.Peters

CHS: Study phenotyping: B.M.P. Genotyping or analysis: J.C.B., K.R., K.D.T. Study PI: B.M.P. AGES-Reykjavik: Genotyping or analysis: A.V.S. Study PI: V.Gudnason, T.B.H., L.J.L.

ERF: Study phenotyping: C.M.v.D., B.A.O. Genotyping or analysis: N.A. Study PI: C.M.v.D., B.A.O. NESDA: Study phenotyping: B.W.J.H.P. Genotyping or analysis: I.M.N., Y.M. Study PI: H.Snieder, B.W.J.H.P.

YFS: Study phenotyping: T.L., M.K., O.T.R. Genotyping or analysis: T.L., L.P.L., M.K., O.T.R. Study PI: T.L., M.K., O.T.R.

EPIC: Genotyping or analysis: N.J.W. Study PI: J.H.Z.

ASPS: Study phenotyping: R.Schmidt Genotyping or analysis: H.Schmidt, E.H., Y.S., R.Schmidt Study

PI: H.Schmidt, R.Schmidt

ORCADES: Study phenotyping: J.F.W., H.C., S.W. Genotyping or analysis: J.F.W., P.K.J., S.W. Study PI: J.F.W.

FINRISK (COROGENE_CTRL): Study phenotyping: P.J. Genotyping or analysis: K.K., A.P.S. Study PI: M.P., P.J.

INGI-VB: Study phenotyping: C.F.S. Genotyping or analysis: M.T., C.M.B., C.F.S. Study PI: D.T. FINRISK_PREDICT_CVD: Study phenotyping: V.S., A.S.H. Study PI: V.S., A.Palotie, S.R.

TRAILS: Study phenotyping: H.R. Genotyping or analysis: P.J.v.d.M. Study PI: C.A.H., A.J.O.

PROCARDIS: Study phenotyping: A.G. Genotyping or analysis: A.G. Study PI: H.W., M.Farrall

HABC: Study phenotyping: Y.Liu, T.B.H. Genotyping or analysis: M.A.N. Study PI: Y.Liu, T.B.H.

KORA S3: Study phenotyping: C.G. Genotyping or analysis: S.S., C.G., E.O. Study PI: M.Laan

INGI-FVG: Genotyping or analysis: D.V., M.Brumat, M.Cocca Study PI: P.G.

Fenland: Study phenotyping: R.A.S., J.a.L., C.L., N.J.W. Genotyping or analysis: R.A.S., J.a.L., C.L., N.J.W. Study PI: R.A.S., C.L., N.J.W.

MICROS: Genotyping or analysis: A.A.H., F.D.G.M., A.S.P. Study PI: F.D.G.M., P.P.P.

HTO: Study phenotyping: B.D.K. Genotyping or analysis: B.D.K., K.L.A., C.Mamasoula Study PI: B.D.K., H.J.C.

MIGEN: Study phenotyping: R.E., J.Marrugat, S.Kathiresan, D.S. Genotyping or analysis: R.E., S.Kathiresan, D.S. Study PI: S.Kathiresan

ULSAM: Study phenotyping: V.Giedraitis, E.I. Genotyping or analysis: A.P.M., A.Mahajan Study PI: A.P.M., V.Giedraitis, E.I.

Cilento study: Study phenotyping: R.Sorice Genotyping or analysis: D.R., T.Nutile Study PI: M.Ciullo LBC1936: Study phenotyping: I.J.D., A.J.G. Genotyping or analysis: L.M.L., G.D., A.J.G. Study PI: I.J.D.

H2000_CTRL: Study phenotyping: T.Niiranen Study PI: P.K., A.Jula, S.Koskinen

NSPHS: Genotyping or analysis: S.E., Å.J. Study PI: U.G.

FUSION: Genotyping or analysis: A.U.J. Study PI: J.T., M.Boehnke, F.C.

GRAPHIC: Study phenotyping: N.J.S., P.S.B., M.D.T. Genotyping or analysis: C.P.N., P.S.B., M.D.T. Study PI: N.J.S.

CROATIA_Vis: Study phenotyping: I.R. Genotyping or analysis: V.V., J.E.H. Study PI: V.V., I.R.

PIVUS: Study phenotyping: L.Lind, J.S. Genotyping or analysis: C.M.L., A.Mahajan Study PI: C.M.L., L.Lind, J.S.

LOLIPOP: Study phenotyping: J.S.K., J.C.C. Genotyping or analysis: J.S.K., W.Z., J.C.C., B.L. Study PI: J.S.K.,J.C.C.

CROATIA_Korcula: Genotyping or analysis: C.H., J.Marten Study PI: C.H., A.F.W.

INGI-CARL: Study phenotyping: G.G. Genotyping or analysis: I.G., A.Morgan, A.R.

LBC1921: Study phenotyping: J.M.S., A.Pattie Genotyping or analysis: J.M.S., S.E.H., D.C.M.L., A.Pattie Study PI: J.M.S.

CROATIA_SPLIT: Study phenotyping: O.P., I.K. Genotyping or analysis: O.P., T.Z. Study PI: O.P. BioMe (formerly IPM): Genotyping or analysis: Y.Lu Study PI: R.J.F.L., E.P.B.

### Replication

UKB-BP: Genotyping or analysis: H.R.W., M.R.B., C.P.C., E.E., H.G., B.M., M.R., I.T. Study PI: P.E., M.J.C.

GoDARTS: Study phenotyping: C.N.A.P., A.S.F.D. Genotyping or analysis: C.N.A.P., N.Shah Study PI: C.N.A.P., A.D.M.

Lifelines: Study phenotyping: M.H.d.B. Genotyping or analysis: M.S. Study PI: P.v.d.H.

TwinsUK: Study phenotyping: C.Menni Genotyping or analysis: M.M., C.Menni Study PI: T.D.S.

Airwave Health Monitoring Study: Genotyping or analysis: A.C.V., E.E., H.G., I.T. Study PI: E.E.

The UK Household Longitudinal Study (UKHLS): Genotyping or analysis: B.P.P. Study PI: E.Z. Generation Scotland (GS:SFHS): Study phenotyping: S.P. Genotyping or analysis: C.H., A.Campbell JUPITER: Study phenotyping: P.M.R. Genotyping or analysis: D.I.C., L.M.R., F.G., P.M.R. Study PI: D.I.C., P.M.R.

NEO: Study phenotyping: R.d.M. Genotyping or analysis: D.O.M.K., R.L.G. Study PI: R.d.M.

Three City-Dijon: Study phenotyping: S.D., C.T. Genotyping or analysis: G.C. Study PI: S.D., C.T. ASCOT-UK: Study phenotyping: P.S., N.P. Genotyping or analysis: P.B.M., H.R.W. Study PI: P.B.M., P.S., N.P., M.J.C.

ASCOT-SC: Study phenotyping: S.Thom, M.J.C. Genotyping or analysis: D.C.S., A.S., H.R.W., P.B.M. Study PI: S.Thom, M.J.C., P.B.M.

Hunter Community Study: Study phenotyping: R.Scott Genotyping or analysis: C.O., E.G.H. Study PI: A.John

GAPP: Study phenotyping: D.C. Genotyping or analysis: D.C., S.Thériault, G.P. Study PI: D.C. BRIGHT: Study phenotyping: M.Brown, J.C. Genotyping or analysis: M.Farrall, P.B.M., H.R.W. Study PI: M.Brown, J.C., M.Farrall, P.B.M., M.J.C.

### Resources for secondary analyses

eQTL NESDA NTR: Design of secondary analysis: R. Jansen Computation of secondary analysis: D.I.B., R.Jansen, B.W.J.H.P. Study PI: D.I.B., B.W.J.H.P.

eQTL kidney: Study phenotyping: J.J.D., M.A.S. Genotyping or analysis: PJ.v.d.M. Study PI: H.Snieder

eQTL BIOS: Design of secondary analysis: R.Jansen Computation of secondary analysis: R.Jansen Study PI: R.Jansen

SABRe: Study phenotyping: Y.D., P.J.M., Q.T.N. Genotyping or analysis: R.Joehanes Design of secondary analysis: D.L. Study PI: D.L.

### ICBP-Steering Committee

G.A., M.J.C., A.Chakravarti, D.I.C., G.B.E., P.E., T.F., M.R.J., A.D.J., M.Larson, D.L., A.P.M., P.B.M., C.N.C., P.O.R., W.P., B.M.P., K.R., A.V.S., H.Snieder, M.D.T., C.M.v.D., L.V.W., H.R.W.

## ACKNOWLEDGMENTS

This research used the ALICE and SPECTRE High Performance Computing Facilities at the University of Leicester. G.B.E is supported by Geneva University Hospitals, Geneva University, de Reuter Foundation, the Swiss National Foundation project FN 33CM30-124087, and the “Fondation pour Recherches Médicales”, Geneva.

### Airwave

We thank all participants of the Airwave Health Monitoring Study. The study is funded by the UK Home Office, (Grant number 780-TETRA) with additional support from the National Institute for Health Research Imperial College Health Care NHS Trust and Imperial College Biomedical Research Centre.

### ARIC

The Atherosclerosis Risk in Communities Study is carried out as a collaborative study supported by National Heart, Lung, and Blood Institute contracts (HHSN268201100005C, HHSN268201100006C, HHSN268201100007C, HHSN268201100008C, HHSN268201100009C, HHSN268201100010C, HHSN268201100011C, and HHSN268201100012C), R01HL087641, R01HL59367 and R01HL086694; National Human Genome Research Institute contract U01HG004402; and National Institutes of Health contract HHSN268200625226C. Funding support for the Genetic Epidemiology of Causal Variants Across the Life Course (CALiCo) program was provided through the NHGRI PAGE program (U01 HG007416). The authors thank the staff and participants of the ARIC study for their important contributions. The authors thank the staff and participants of the ARIC study for their important contributions.

### ASCOT

This work was supported by Pfizer, New York, NY, USA, for the ASCOT study and the collection of the ASCOT DNA repository; by Servier Research Group, Paris, France; and by Leo Laboratories, Copenhagen, Denmark. We thank all ASCOT trial participants, physicians, nurses, and practices in the participating countries for their important contribution to the study. In particular we thank Clare Muckian and David Toomey for their help in DNA extraction, storage, and handling. This work forms part of the research programme of the NIHR Cardiovascular Biomedical Research Unit at Barts

### ASPS

The research reported in this article was funded by the Austrian Science Fond (FWF) grant number P20545-P05 and P13180. The Medical University of Graz supports the databank of the ASPS. The authors thank the staff and the participants of the ASPS for their valuable contributions. The authors thank Birgit Reinhart for her long-term administrative commitment and Ing Johann Semmler for the technical assistance at creating the DNA bank.

### BRIGHT

This work was supported by the Medical Research Council of Great Britain (grant number G9521010D); and by the British Heart Foundation (grant number PG/02/128). The BRIGHT study is extremely grateful to all the patients who participated in the study and the BRIGHT nursing team. This work forms part of the research programme of the NIHR Cardiovascular Biomedical Research Unit at Barts.

### B58C

We acknowledge use of phenotype and genotype data from the British 1958 Birth Cohort DNA collection, funded by the Medical Research Council grant G0000934 and the Wellcome Trust grant 068545/Z/02. Genotyping for the B58C-WTCCC subset was funded by the Wellcome Trust grant 076113/B/04/Z. The B58C-T1DGC genotyping utilized resources provided by the Type 1 Diabetes Genetics Consortium, a collaborative clinical study sponsored by the National Institute of Diabetes and Digestive and Kidney Diseases (NIDDK), National Institute of Allergy and Infectious Diseases (NIAID), National Human Genome Research Institute (NHGRI), National Institute of Child Health and Human Development (NICHD), and Juvenile Diabetes Research Foundation International (JDRF) and supported by U01 DK062418. B58C-T1DGC GWAS data were deposited by the Diabetes and Inflammation Laboratory, Cambridge Institute for Medical Research (CIMR), University of Cambridge, which is funded by Juvenile Diabetes Research Foundation International, the Wellcome Trust and the National Institute for Health Research Cambridge Biomedical Research Centre; the CIMR is in receipt of a Wellcome Trust Strategic Award (079895). The B58C-GABRIEL genotyping was supported by a contract from the European Commission Framework Programme 6 (018996) and grants from the French Ministry of Research.

### CHS

This CHS research was supported by NHLBI contracts HHSN268201200036C, HHSN268200800007C, N01HC55222, N01HC85079, N01HC85080, N01HC85081, N01HC85082, N01HC85083, N01HC85086, HHSN268200960009C; and NHLBI grants U01HL080295, R01HL087652, R01HL105756, R01HL103612, R01HL120393, and R01HL130114 with additional contribution from the National Institute of Neurological Disorders and Stroke (NINDS). Additional support was provided through R01AG023629 from the National Institute on Aging (NIA). A full list of principal CHS investigators and institutions can be found at CHS-NHLBI.org. The provision of genotyping data was supported in part by the National Center for Advancing Translational Sciences, CTSI grant UL1TR000124, and the National Institute of Diabetes and Digestive and Kidney Disease Diabetes Research Center (DRC) grant DK063491 to the Southern California Diabetes Endocrinology Research Center. The content is solely the responsibility of the authors and does not necessarily represent the official views of the National Institutes of Health.

### Cilento study

The Cilento study was supported by the Italian Ministry of Education Universities and Research (Interomics Flagship Project, PON03PE_00060_7), FP6 (Vasoplus-037254), the Assessorato Ricerca Regione Campania, the Fondazione con il SUD (2011-PDR-13), and the Istituto Banco di Napoli - Fondazione to MC. We address special thanks to the populations of Cilento for their participation in the study.

### COLAUS

The CoLaus study was and is supported by research grants from GlaxoSmithKline (GSK), the Faculty of Biology and Medicine of Lausanne, and the Swiss National Science Foundation (grants 3200B0-105993, 3200B0-118308, 33CSCO-122661, and 33CS30-139468). We thank all participants, involved physicians and study nurses to the CoLaus cohort.

### COROGENE_CTRL

This study has been funded by the Academy of Finland (grant numbers 139635, 129494, 118065, 129322, 250207), the Orion-Farmos Research Foundation, the Finnish Foundation for Cardiovascular Research, and the Sigrid Jusélius Foundation. We are grateful for the THL DNA laboratory for its skillful work to produce the DNA samples used in this study. We thank the Sanger Institute genotyping facilities for genotyping the samples.

### CROATIA Studies

The CROATIA-Vis, CROATIA-Korcula and CROATIA-Split studies in the Croatian islands of Vis and Korcula and mainland city of Split were supported by grants from the Medical Research Council (UK); the Ministry of Science, Education, and Sport of the Republic of Croatia (grant number 216-1080315-0302); the European Union framework program 6 European Special Populations Research Network project (contract LSHG-CT-2006-018947), the European Union framework program 7 project BBMRI-LPC (FP7 313010) and the Croatian Science Foundation (grant 8875). The CROATIA studies would like to acknowledge the invaluable contributions of the recruitment teams (including those from the Institute of Anthropological Research in Zagreb) in Vis, Korcula and Split, the administrative teams in Croatia and Edinburgh, and the people of Vis, Korcula and Split. SNP genotyping of the CROATIA-Vis samples was carried out by the Genetics Core Laboratory at the Wellcome Trust Clinical Research Facility, WGH, Edinburgh, Scotland. SNP genotyping for CROATIA-Korcula was performed by Helmholtz ZentrumMünchen, GmbH, Neuherberg, Germany. The SNP genotyping for the CROATIA-Split cohort was performed by AROS Applied Biotechnology, Aarhus, Denmark.

### ERF

The ERF study as a part of EUROSPAN (European Special Populations Research Network) was supported by European Commission FP6 STRP grant number 018947 (LSHG-CT-2006-01947) and also received funding from the European Community's Seventh Framework Programme (FP7/2007-2013)/grant agreement HEALTH-F4-2007-201413 by the European Commission under the programme "Quality of Life and Management of the Living Resources" of 5th Framework Programme (no. QLG2-CT-2002-01254). High-throughput analysis of the ERF data was supported by a joint grant from the Netherlands Organization for Scientific Research and the Russian Foundation for Basic Research (NWO-RFBR 047.017.043). Exome sequencing analysis in ERF was supported by the ZonMw grant (project 91111025). Najaf Amin is supported by the Netherlands Brain Foundation (project number F2013(1)-28). We are grateful to all study participants and their relatives, general practitioners and neurologists for their contributions and to P. Veraart for her help in genealogy, J. Vergeer for the supervision of the laboratory work and P. Snijders for his help in data collection.

### Fenland

JAL, CL, RAS and NJW acknowledge support from the Medical Research Council (MC_U106179471 and MC_UU_12015/1) The Fenland Study is funded by the Wellcome Trust and the Medical Research Council (MC_U106179471). We are grateful to all the volunteers for their time and help, and to the General Practitioners and practice staff for assistance with recruitment. We thank the Fenland Study Investigators, Fenland Study Co-ordination team and the Epidemiology Field, Data and Laboratory teams. We further acknowledge support from the Medical research council (MC_UU_12015/1).

### FHS

The National Heart, Lung and Blood Institute’s Framingham Heart Study is supported by contract N01-HC-25195

### FINRISK_PREDICT_CVD

This study has been funded by the Academy of Finland (grant numbers 139635, 129494, 118065, 129322, 250207, 269517), the Orion-Farmos Research Foundation, the Finnish Foundation for Cardiovascular Research, and the Sigrid Jusélius Foundation. We are grateful for the THL DNA laboratory for its skillful work to produce the DNA samples used in this study. We thank the Sanger Institute genotyping facilities for genotyping the samples.

### FUSION

Support for FUSION was provided by NIH grants R01-DK062370 (to M.B.) and intramural project number ZIA-HG000024 (to F.S.C.). Genome-wide genotyping was conducted by the Johns Hopkins University Genetic Resources Core Facility SNP Center at the Center for Inherited Disease Research (CIDR), with support from CIDR NIH contract no. N01-HG-65403.

### GAPP study

The GAPP study was supported by the Liechtenstein Government, the Swiss National Science Foundation, the Swiss Heart Foundation, the Swiss Society of Hypertension, the University of Basel, the University Hospital Basel, the Hanela Foundation, Schiller AG and Novartis.

### GS:SFHS

Generation Scotland received core support from the Chief Scientist Office of the Scottish Government Health Directorates [CZD/16/6] and the Scottish Funding Council [HR03006]. Genotyping of the GS:SFHS samples was carried out by the Genetics Core Laboratory at the Wellcome Trust Clinical Research Facility, Edinburgh, Scotland and was funded by the Medical Research Council UK and the Wellcome Trust (Wellcome Trust Strategic Award “STratifying Resilience and Depression Longitudinally” (STRADL) Reference 104036/Z/14/Z). Ethics approval for the study was given by the NHS Tayside committee on research ethics (reference 05/S1401/89). We are grateful to all the families who took part, the general practitioners and the Scottish School of Primary Care for their help in recruiting them, and the whole Generation Scotland team, which includes interviewers, computer and laboratory technicians, clerical workers, research scientists, volunteers, managers, receptionists, healthcare assistants and nurses.

### GoDARTS

GoDARTS was funded by The Wellcome Trust (072960/Z/03/Z, 084726/Z/08/Z, 084727/Z/08/Z, 085475/Z/08/Z, 085475/B/08/Z) and as part of the EU IMI-SUMMIT program. We acknowledge the support of the Health Informatics Centre, University of Dundee for managing and supplying the anonymised data and NHS Tayside, the original data owner. We are grateful to all the participants who took part in the Go-DARTS study, to the general practitioners, to the Scottish School of Primary Care for their help in recruiting the participants, and to the whole team, which includes interviewers, computer and laboratory technicians, clerical workers, research scientists, volunteers, managers, receptionists, and nurses.

### GRAPHIC

The GRAPHIC Study was funded by the British Heart Foundation (BHF/RG/2000004). CPN and NJS are supported by the British Heart Foundation and NJS is a NIHR Senior Investigator This work falls under the portfolio of research supported by the NIHR Leicester Cardiovascular Biomedical Research Unit.

### H2000

The Health 2000 Study was funded by the National Institute for Health and Welfare (THL), the Finnish Centre for Pensions (ETK), the Social Insurance Institution of Finland (KELA), the Local Government Pensions Institution (KEVA) and other organizations listed on the website of the survey (http://www.terveys2000.fi). We are grateful for the THL DNA laboratory for its skillful work to produce the DNA samples used in this study. We thank the Sanger Institute genotyping facilities for genotyping the GenMets subcohort.

### HABC

The Health ABC Study was supported by NIA contracts N01AG62101, N01AG62103, and N01AG62106 and, in part, by the NIA Intramural Research Program. The genome-wide association study was funded by NIA grant 1R01AG032098-01A1 to Wake Forest University Health Sciences and genotyping services were provided by the Center for Inherited Disease Research (CIDR). CIDR is fully funded through a federal contract from the National Institutes of Health to The Johns Hopkins University, contract number HHSN268200782096C. This study utilized the high-performance computational capabilities of the Biowulf Linux cluster at the National Institutes of Health, Bethesda, Md. (http://biowulf.nih.gov).

### HTO

The study was funded by the Wellcome Trust, Medical Research Council and British Heart Foundation We thank all the families who participated in the study

### INGI-VB

The INGI-Val Borbera population is a collection of 1,664 genotyped samples collected in the Val Borbera Valley, a geographically isolated valley located within the Appennine Mountains in Northwest Italy. The valley is inhabited by about 3,000 descendants from the original population, living in 7 villages along the valley and in the mountains. Participants were healthy people 18-102 years of age that had at least one grandfather living in the valley. The study plan and the informed consent form were reviewed and approved by the institutional review boards of San Raffaele Hospital in Milan. The research was supported by funds from Compagnia di San Paolo, Torino, Italy; Fondazione Cariplo, Italy and Ministry of Health, Ricerca Finalizzata 2008 and CCM 2010, PRIN 2009 and Telethon, Italy to DT. The funders had no role in study design, data collection and analysis, decision to publish, or preparation of the manuscript. We thank the inhabitants of the VB that made this study possible, the local administrations, the MD of the San Raffaele Hospital and Prof Clara Camaschella for clinical data collection. We also thank Fiammetta Viganò for technical help, Corrado Masciullo and Massimiliano Cocca for building and maintaining the analysis platform.

### INGI-CARL

Italian Ministry of Health RF2010 to Paolo Gasparini, RC2008 to Paolo Gasparini

### INGI-FVG

Italian Ministry of Health RF2010 to Paolo Gasparini, RC2008 to Paolo Gasparini

### JUPITER

Genetic analysis in the JUPITER trial was funded by a grant from AstraZeneca (DIC and PMR, Co-Pis).

### KORA S3

KORA S3 500K blood pressure project was supported by Estonian Research Council, grant IUT34-12 (for Maris Laan). The KORA Augsburg studies have been financed by the Helmholtz Zentrum Mu nchen, German Research Center for Environmental Health, Neuherberg, Germany and supported by grants from the German Federal Ministry of Education and Research (BMBF). The KORA study group consists of H-E. Wichmann (speaker), A. Peters, C. Meisinger, T. Illig, R. Holle, J. John and co-workers, who are responsible for the design and conduct of the KORA studies. Part of this work was financed by the German National Genome Research Network (NGFN-2 and NGFNPlus:01GS0823) and supported within the Munich Center of Health Sciences (MC Health) as part of LMUinnovativ.

### LBC1921

Phenotype collection in the Lothian Birth Cohort 1921 was supported by the UK’s Biotechnology and Biological Sciences Research Council (BBSRC), The Royal Society, and The Chief Scientist Office of the Scottish Government. Genotyping was funded by the BBSRC. The work was undertaken by The University of Edinburgh Centre for Cognitive Ageing and Cognitive Epidemiology, part of the cross council Lifelong Health and Wellbeing Initiative (MR/K026992/1). Funding from the BBSRC and Medical Research Council (MRC) is gratefully acknowledged. We thank the Lothian Birth Cohort 1921 participants and team members who contributed to these studies.

### LBC1936

Phenotype collection in the Lothian Birth Cohort 1936 was supported by Age UK (The Disconnected Mind project). Genotyping was funded by the BBSRC. The work was undertaken by The University of Edinburgh Centre for Cognitive Ageing and Cognitive Epidemiology, part of the cross council Lifelong Health and Wellbeing Initiative (MR/K026992/1). Funding from the BBSRC and Medical Research Council (MRC) is gratefully acknowledged. We thank the Lothian Birth Cohort 1936 participants and team members who contributed to these studies.

### Lifelines Cohort Study

The LifeLines Cohort Study, and generation and management of GWAS genotype data for the LifeLines Cohort Study is supported by the Netherlands Organization of Scientific Research NWO (grant 175.010.2007.006), the Economic Structure Enhancing Fund (FES) of the Dutch government, the Ministry of Economic Affairs, the Ministry of Education, Culture and Science, the Ministry for Health, Welfare and Sports, the Northern Netherlands Collaboration of Provinces (SNN), the Province of Groningen, University Medical Center Groningen, the University of Groningen, Dutch Kidney Foundation and Dutch Diabetes Research Foundation. The authors wish to acknowledge the services of the Lifelines Cohort Study, the contributing research centers delivering data to Lifelines, and all the study participants.

### LOLIPOP

The LOLIPOP study is funded by the British Heart Foundation (SP/04/002), the Medical Research Council (G0601966, G0700931), the Wellcome Trust (084723/Z/08/Z), the NIHR (RP-PG-0407-10371), European Union FP7 (EpiMigrant, 279143) and Action on Hearing Loss (G51). The LOLIPOP study is supported by the National Institute for Health Research (NIHR) Comprehensive Biomedical Research Centre Imperial College Healthcare NHS Trust. The work was carried out in part at the NIHR/Wellcome Trust Imperial Clinical Research Facility. We thank the participants and research staff who made the study possible.

### MESA

This research was supported by the Multi-Ethnic Study of Atherosclerosis (MESA) contracts N01-HC-95159, N01-HC-95160, N01-HC-95161, N01-HC-95162, N01-HC-95163, N01-HC-95164, N01- HC-95165, N01-HC-95166, N01-HC-95167, N01-HC-95168, N01-HC-95169 and by grants UL1-TR-000040 and UL1-RR-025005 from NCRR. Funding for MESA SHARe genotyping was provided by NHLBI Contract N02-HL-6-4278. The provision of genotyping data was supported in part by the National Center for Advancing Translational Sciences, CTSI grant UL1TR000124, and the National Institute of Diabetes and Digestive and Kidney Disease Diabetes Research Center (DRC) grant DK063491 to the Southern California Diabetes Endocrinology Research Center.

### MICROS

The MICROS study was supported by the Ministry of Health and Department of Innovation, Research and University of the Autonomous Province of Bolzano, the South Tyrolean Sparkasse Foundation, and the European Union framework program 6 EUROSPAN project (contract no. LSHG-CT-2006-018947) For the MICROS study, we thank the primary care practitioners Raffaela Stocker, Stefan Waldner, Toni Pizzecco, Josef Plangger, Ugo Marcadent, and the personnel of the Hospital of Silandro (Department of Laboratory Medicine) for their participation and collaboration in the research project.

### NEO

The NEO study is supported by the participating Departments, the Division and the Board of Directors of the Leiden University Medical Center, and by the Leiden University, Research Profile Area Vascular and Regenerative Medicine. Dennis Mook-Kanamori is supported by Dutch Science Organization (ZonMW-VENI Grant 916.14.023). The authors of the NEO study thank all individuals who participated in the Netherlands Epidemiology in Obesity study, all participating general practitioners for inviting eligible participants and all research nurses for collection of the data. We thank the NEO study group, Pat van Beelen, Petra Noordijk and Ingeborg de Jonge for the coordination, lab and data management of the NEO study. The genotyping in the NEO study was supported by the Centre National de Génotypage (Paris, France), headed by Jean-Francois Deleuze.

### NESDA

Funding was obtained from the Netherlands Organization for Scientific Research (Geestkracht program grant 10-000-1002); the Center for Medical Systems Biology (CSMB, NOW Genomics), Biobanking and Biomolecular Resources Research Infrastructure (BBMRI-NL), VU University’s Institutes for Health and Care Research (EMGO+) and Neuroscience Campus Amsterdam, University Medical Center Groningen, Leiden University Medical Center, National Institutes of Health (NIH, R01D0042157-01A, MH081802, Grand Opportunity grants 1RC2 MH089951 and 1RC2 MH089995). Part of the genotyping and analyses were funded by the Genetic Association Information Network (GAIN) of the Foundation for the National Institutes of Health. Computing was supported by BiG Grid, the Dutch e-Science Grid, which is financially supported by NWO.

### NSPHS

The Northern Swedish Population Health Study (NSPHS) was funded by the Swedish Medical Research Council (Project Number K2007-66X-20270-01-3, 2011-5252, 2012-2884 and 2011-2354), the Foundation for Strategic Research (SSF). NSPHS as part of EUROSPAN (European Special Populations Research Network) was also supported by the European Commission FP6 STRP grant number 01947 (LSHG-CT-2006-01947). This work has also been supported by the Swedish Society for Medical Research (SSMF), and the Swedish Medical Research Council (#2015-03327) We are grateful for the contribution of district nurse Svea Hennix for data collection and Inger Jonasson for logistics and coordination of the health survey. We also thank all the participants from the community for their interest and willingness to contribute to this study.

### NTR

Funding was obtained from the Netherlands Organization for Scientific Research (NWO) and The Netherlands Organisation for Health Research and Development (ZonMW) grants 904-61-090, 985-10-002, 904-61-193,480-04-004, 400-05-717, Addiction-31160008, Middelgroot-911-09-032, Spinozapremie 56-464-14192, Biobanking and Biomolecular Resources Research Infrastructure (BBMRI-NL, 184.021.007); the Netherlands Heart Foundation grants 86.083 and 88.042 and 90.313; the VU Institute for Health and Care Research (EMGO+); the European Community’s Seventh Framework Program (FP7/2007-2013), ENGAGE (HEALTH-F4-2007-201413); the European Research Council (ERC Advanced, 230374), the Rutgers University Cell and DNA Repository (NIMH U24 MH068457-06), the Avera Institute, Sioux Falls, South Dakota (USA) and the National Institutes of Health (NIH, R01D0042157-0ΙΑ, MH081802; Grand Opportunity grant 1RC2 MH089951). Part of the genotyping and analyses were funded by the Genetic Association Information Network (GAIN) of the Foundation for the National Institutes of Health. Computing was supported by BiG Grid, the Dutch e-Science Grid, which is financially supported by NWO.”

### ORCADES

ORCADES was supported by the Chief Scientist Office of the Scottish Government, the Royal Society, the MRC Human Genetics Unit, Arthritis Research UK and the European Union framework program 6 EUROSPAN project (contract no. LSHG-CT-2006-018947). DNA extractions were performed at the Wellcome Trust Clinical Research Facility in Edinburgh. We would like to acknowledge the invaluable contributions of Lorraine Anderson and the research nurses in Orkney, the administrative team in Edinburgh and the people of Orkney.

### PIVUS

This project was supported by Knut and Alice Wallenberg Foundation (Wallenberg Academy Fellow), European Research Council (ERC Starting Grant), Swedish Diabetes Foundation (grant no. 2013-024), Swedish Research Council (grant no. 2012-1397), and Swedish Heart-Lung Foundation (20120197). The computations were performed on resources provided by SNIC through Uppsala Multidisciplinary Center for Advanced Computational Science (UPPMAX) under Project b2011036. Genetic data analysis was funded by the Wellcome Trust under awards WT098017 and WT090532. We thank the SNP&SEQ Technology Platform in Uppsala (http://www.genotyping.se) for excellent genotyping.

### PROCARDIS

PROCARDIS was supported by the European Community Sixth Framework Program (LSHM-CT- 2007-037273), AstraZeneca, the British Heart Foundation, the Swedish Research Council, the Knut and Alice Wallenberg Foundation, the Swedish Heart-Lung Foundation, the Torsten and Ragnar Söderberg Foundation, the Strategic Cardiovascular Program of Karolinska Institutet and Stockholm County Council, the Foundation for Strategic Research and the Stockholm County Council (560283). M.F and H.W acknowledge the support of the Wellcome Trust core award (090532/Z/09/Z) and M.F, H.W, the BHF Centre of Research Excellence (RE/13/1/30181). A.G, H.W acknowledge European Union Seventh Framework Programme FP7/2007-2013 under grant agreement no. HEALTH-F2-2013-601456 (CVGenes@Target) & and A.G, the Wellcome Trust Institutional strategic support fund. PROCARDIS was supported by the European Community Sixth Framework Program (LSHM-CT- 2007-037273), AstraZeneca, the British Heart Foundation, the Swedish Research Council, the Knut and Alice Wallenberg Foundation, the Swedish Heart-Lung Foundation, the Torsten and Ragnar Söderberg Foundation, the Strategic Cardiovascular Program of Karolinska Institutet and Stockholm County Council, the Foundation for Strategic Research and the Stockholm County Council (560283). M.F and H.W acknowledge the support of the Wellcome Trust core award (090532/Z/09/Z) and M.F, H.W, the BHF Centre of Research Excellence (RE/13/1/30181). A.G, H.W acknowledge European Union Seventh Framework Programme FP7/2007-2013 under grant agreement no. HEALTH-F2-2013-601456 (CVGenes@Target) & and A.G, the Wellcome Trust Institutional strategic support fund.

### PROSPER

The PROSPER study was supported by an investigator initiated grant obtained from Bristol-Myers Squibb. Prof. Dr. J. W. Jukema is an Established Clinical Investigator of the Netherlands Heart Foundation (grant 2001 D 032). Support for genotyping was provided by the seventh framework program of the European commission (grant 223004) and by the Netherlands Genomics Initiative (Netherlands Consortium for Healthy Aging grant 050-060-810).

### RS

The generation and management of GWAS genotype data for the Rotterdam Study (RS I, RS II, RS III) was executed by the Human Genotyping Facility of the Genetic Laboratory of the Department of Internal Medicine, Erasmus MC, Rotterdam, The Netherlands. The GWAS datasets are supported by the Netherlands Organisation of Scientific Research NWO Investments (nr. 175.010.2005.011, 91103-012), the Genetic Laboratory of the Department of Internal Medicine, Erasmus MC, the Research Institute for Diseases in the Elderly (014-93-015; RIDE2), the Netherlands Genomics Initiative (NGI)/Netherlands Organisation for Scientific Research (NWO) Netherlands Consortium for Healthy Aging (NCHA), project nr. 050-060-810. The Rotterdam Study is funded by Erasmus Medical Center and Erasmus University, Rotterdam, Netherlands Organization for the Health Research and Development (ZonMw), the Research Institute for Diseases in the Elderly (RIDE), the Ministry of Education, Culture and Science, the Ministry for Health, Welfare and Sports, the European Commission (DG XII), and the Municipality of Rotterdam. We thank Pascal Arp, Mila Jhamai, Marijn Verkerk, Lizbeth Herrera and Marjolein Peters, MSc, and Carolina Medina-Gomez, MSc, for their help in creating the GWAS database, and Karol Estrada, PhD, Yurii Aulchenko, PhD, and Carolina Medina-Gomez, MSc, for the creation and analysis of imputed data.The authors are grateful to the study participants, the staff from the Rotterdam Study and the participating general practitioners and pharmacists.

### SHIP

SHIP is part of the Community Medicine Research net of the University of Greifswald, Germany, which is funded by the Federal Ministry of Education and Research (grants no. 01ZZ9603, 01ZZ0103, and 01ZZ0403), the Ministry of Cultural Affairs as well as the Social Ministry of the Federal State of Mecklenburg-West Pomerania, and the network ‘Greifswald Approach to Individualized Medicine (GANI_MED)’ funded by the Federal Ministry of Education and Research (grant 03IS2061A). Genome-wide data have been supported by the Federal Ministry of Education and Research (grant no. 03ZIK012) and a joint grant from Siemens Healthcare, Erlangen, Germany and the Federal State of Mecklenburg- West Pomerania. The University of Greifswald is a member of the Caché Campus program of the InterSystems GmbH.

### Three City- Dijon

The 3-City Study is conducted under a partnership agreement among the Institut National de la Santé et de la Recherche Médicale (INSERM), the University of Bordeaux, and Sanofi-Aventis. The Fondation pour la Recherche Médicale funded the preparation and initiation of the study. The 3C Study is also supported by the Caisse Nationale Maladie des Travailleurs Salariés, Direction Générale de la Santé, Mutuelle Générale de l’Education Nationale (MGEN), Institut de la Longévité, Conseils Régionaux of Aquitaine and Bourgogne, Fondation de France, and Ministry of Research-INSERM Programme “Cohortes et collections de données biologiques.” This work was supported by the National Foundation for Alzheimer’s Disease and Related Disorders, the Institut Pasteur de Lille, the Centre National de Génotypage and the LABEX (Laboratory of Excellence program investment for the future) DISTALZ - Development of Innovative Strategies for a Transdisciplinary approach to ALZheimer’s disease. Ganesh Chauhan, Christophe Tzourio and Stéphanie Debette are supported by a grant from the Fondation Leducq. We thank Philippe Amouyel and the UMR1167 Inserm Univ Lille Institut Pasteur de Lille for providing the 3C Dijon cohort SNP replication data funded by a grant from the French National Foundation on Alzheimer’s disease and related disorders.

### UKHLS

This work was funded through generous grants from the Economic & Social Research Council (ES/H029745/1) and the Wellcome Trust (WT098051).

### TRAILS

This research is part of the TRacking Adolescents' Individual Lives Survey (TRAILS). Participating centers of TRAILS include the University Medical Center and University of Groningen, the Erasmus University Medical Center Rotterdam, the University of Utrecht, the Radboud Medical Center Nijmegen, and the Parnassia Bavo group, all in the Netherlands. TRAILS has been financially supported by various grants from the Netherlands Organization for Scientific Research NWO (Medical Research Council program grant GB-MW 940-38-011; ZonMW Brainpower grant 100-001-004; ZonMw Risk Behavior and Dependence grants 60-60600-97-118; ZonMw Culture and Health grant 261-98-710; Social Sciences Council medium-sized investment grants GB-MaGW 480-01-006 and GB-MaGW 480-07-001; Social Sciences Council project grants GB-MaGW 452-04-314 and GB-MaGW 452-06-004; NWO large-sized investment grant 175.010.2003.005; NWO Longitudinal Survey and Panel Funding 481-08-013 and 481-11-001), the Dutch Ministry of Justice (WODC), the European Science Foundation (EuroSTRESS project FP-006), Biobanking and Biomolecular Resources Research Infrastructure BBMRI-NL (CP 32), and the participating universities. Statistical analyses were carried out on the Genetic Cluster Computer (http://www.geneticcluster.org) hosted by SURFsara and financially supported by the Netherlands Scientific Organization (NWO 480-05-003 PI: Posthuma) along with a supplement from the Dutch Brain Foundation and the VU University Amsterdam.

### TwinGene

This project was supported by grants from the Ministry for Higher Education, the Swedish Research Council (M-2005-1112 and 2009-2298), GenomEUtwin (EU/QLRT-2001-01254; QLG2-CT-2002-01254), NIH grant DK U01-066134, Knut and Alice Wallenberg Foundation (Wallenberg Academy Fellow), European Research Council (ERC Starting Grant), Swedish Diabetes Foundation (grant no. 2013-024), Swedish Research Council (grant no. 2012-1397), and Swedish Heart-Lung Foundation (20120197). We thank the SNP&SEQ Technology Platform in Uppsala (http://www.genotyping.se) for excellent genotyping. The computations were performed on resources provided by SNIC through Uppsala Multidisciplinary Center for Advanced Computational Science (UPPMAX) under Project b2011036.

### TwinsUK

The study was funded by the Wellcome Trust; European Community’s Seventh Framework Programme (FP7/2007-2013). The study also receives support from the National Institute for Health Research (NIHR) BioResource Clinical Research Facility and Biomedical Research Centre based at Guy's and St Thomas' NHS Foundation Trust and King's College London (guysbrc-2012-1). We thank the staff from the Genotyping Facilities at the Wellcome Trust Sanger Institute for sample preparation, quality control, and genotyping; Le Centre National de Génotypage, France, for genotyping; Duke University, NC, USA, for genotyping; and the Finnish Institute of Molecular Medicine, Finnish Genome Center, University of Helsinki. Genotyping was also done by CIDR as part of an NEI/NIH project grant

### UK Biobank_Cardiometabolic Consortium

This research has been conducted using the UK Biobank Resource under application number 236. H.R.W., C.P.C and M.R.B. were funded by the National Institutes for Health Research (NIHR) as part of the portfolio of translational research of the NIHR Biomedical Research Unit at Barts MR was funded by the National Institute for Health Research (NIHR) Biomedical Research Unit in Cardiovascular Disease at Barts. MR is recipient from China Scholarship Council (No. 2011632047). B.M. holds an MRC eMedLab Medical Bioinformatics Career Development Fellowship, funded from award MR/L016311/1. PE was funded by the National Institutes for Health Research (NIHR) Imperial College Health Care NHS Trust and Imperial College London Biomedical Research Centre, the UK Medical Research Council and Public Health England as Director of the MRC-PHE Centre for Environment and Health, and the NIHR Health Protection Research Unit on the Health Effects of Environmental Hazards. Some of this work used computing resources provided by the Medical Research Council-funded UK MEDical Bioinformatics partnership programme (UK MED-BIO) (MR/L01632X/1).

### ULSAM

This project was supported by Knut and Alice Wallenberg Foundation (Wallenberg Academy Fellow), European Research Council (ERC Starting Grant), Swedish Diabetes Foundation (grant no. 2013-024), Swedish Research Council (grant no. 2012-1397), and Swedish Heart-Lung Foundation (20120197). The computations were performed on resources provided by SNIC through Uppsala Multidisciplinary Center for Advanced Computational Science (UPPMAX) under Project b2011036. Genotyping was funded by the Wellcome Trust under award WT064890. Analysis of genetic data was funded by the Wellcome Trust under awards WT098017 and WT090532. Andrew P. Morris is a Wellcome Trust Senior Research Fellow in Basic Biomedical Science (WT098017). We thank the SNP&SEQ Technology Platform in Uppsala (http://www.genotyping.se) for excellent genotyping.

### WGHS

The WGHS is supported by the National Heart, Lung, and Blood Institute (HL043851, HL080467, HL09935) and the National Cancer Institute (CA047988 and UM1CA182913) with collaborative scientific support and funding for genotyping provided by Amgen.

### YFS

The Young Finns Study has been financially supported by the Academy of Finland: grants 286284, 134309 (Eye), 126925, 121584, 124282, 129378 (Salve), 117787 (Gendi), and 41071 (Skidi); the Social Insurance Institution of Finland; Kuopio, Tampere and Turku University Hospital Medical Funds (grant X51001); Juho Vainio Foundation; Paavo Nurmi Foundation; Finnish Foundation for Cardiovascular Research; Finnish Cultural Foundation; Tampere Tuberculosis Foundation; Emil Aaltonen Foundation; Yrjö Jahnsson Foundation; Signe and Ane Gyllenberg Foundation; and Diabetes Research Foundation of Finnish Diabetes Association. The expert technical assistance in the statistical analyses by Irina Lisinen is gratefully acknowledged.

